# Extraordinary Self-Sacrifice

**DOI:** 10.1101/178376

**Authors:** D.B. Krupp

## Abstract

Theories of self-sacrifice ordinarily assume that actors will have larger fitness effects on recipients than on themselves. There are, however, conditions in which actors can pay costs that exceed the altruistic benefits they provide or the spiteful costs they impose. In a spatially structured population, I show that such “extraordinary” self-sacrifice evolves when actors use information about kinship and dispersal to maximize inclusive fitness. The result can be described by a simple rule: extraordinary self-sacrifice evolves when the actor’s neighbors are kin and the recipient’s neighbors are not.

---

Inclusive fitness theory distinguishes two kinds of self-sacrifice: altruism and spite. Whereas altruism entails a lifetime fitness cost to the actor (*C* > 0) and a benefit to the recipient (*B* > 0), spite entails a cost to the actor (*C* > 0) and a cost to the recipient (*B* < 0). Both are expected to evolve when they satisfy Hamilton’s rule,

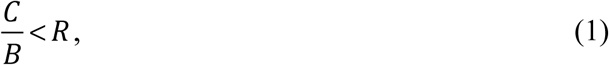

where *R* is the genetic relatedness between the actor and the recipient (*1*-*5*).

It is ordinarily assumed that actors cannot pay costs that exceed their effects on recipients, setting a practical limit of *C*/|*B*| ≤ 1 on the evolution of self-sacrifice. Individuals are thus expected to value their own reproduction and survival more than the reproduction and survival of others. Accordingly, it is difficult to explain the existence of adaptations that exact severe costs on actor fitness, such as irreversible sterility and self-destructive weaponry (*5*, *6*).

Here, I revisit connections between kinship and dispersal made by Hamilton fifty years ago (*7*) and Taylor twenty-five years ago (*8*) to show how self-sacrifice beyond the *C*/|*B*| ≤ 1 limit can evolve. My argument rests on the ability of actors to steer costs away from kin and benefits toward them. Costs and benefits are typically measured in the context of the immediate or “primary” interaction between the actor and recipient. However, these primary effects can have downstream or “secondary” consequences for neighbors, who may also share identical copies of the allele causing the actor’s behavior (*2*-*5*, *8*-*13*). In a Malthusian (i.e. inelastic) population, the primary effects –*C* and *B* cause equal and opposite secondary effects *C* and –*B* on the actor’s and recipient’s neighbors, respectively, who compete for opportunities to breed. If those neighbors are kin, then the secondary effects of a social action will also influence selection (Fig. 1).

**Fig. 1.**
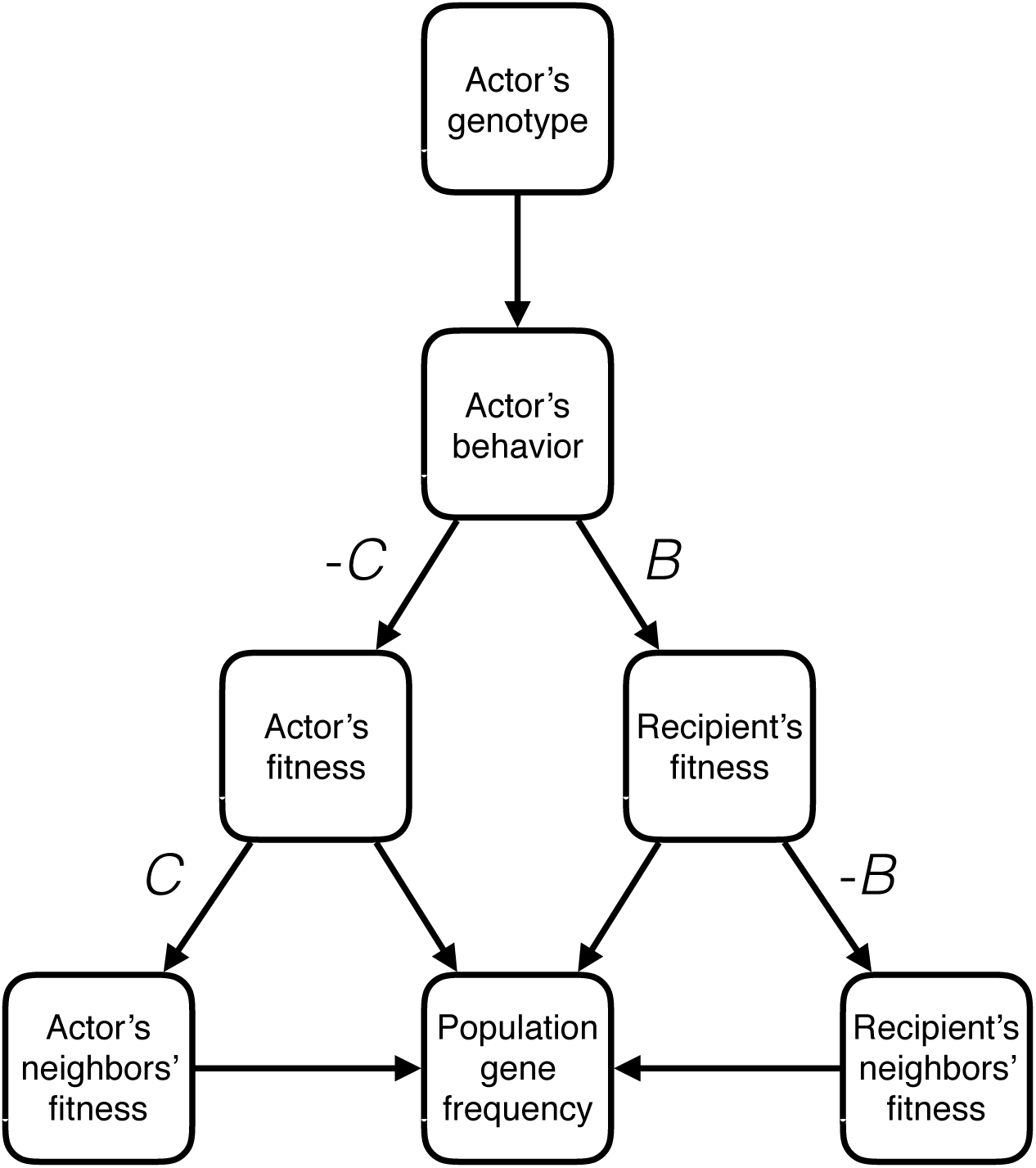
Social evolution in a Malthusian population. A gene causes the actor’s behavior, which has a primary effect –*C* on the actor’s fitness and a primary effect *B* on the recipient’s fitness. However, the action also has a secondary effect *C* on the actor’s neighbors’ fitness and a secondary effect –*B* on the recipient’s neighbors’ fitness. Consequently, both the primary and secondary effects can determine the frequency of the focal allele in the population.

This logic suggests that self-sacrifice depends on the reduction of reproductive competition among kin and an increase of the same among nonkin. The coordination of three variables, in particular, would give rise to this. The first is the probability of the actor’s own dispersal. If the actor remains at home, the secondary benefit of its sacrifice goes to kin; conversely, if the actor disperses, the secondary benefit goes to nonkin. The second is the probability of the recipient’s dispersal. If the recipient remains at home, the secondary effect is conferred upon the actor’s kin; conversely, if the recipient disperses, the secondary effect is conferred upon nonkin. The third is the probability that the actor and the recipient share copies of the focal allele identical by descent, also known as consanguinity. If the two parties are kin, then altruism can evolve, whereas if they are nonkin, then spite can evolve.

I study the effects of these variables in a series of spatially structured, analytical models (*8*, *14*, *15*). I focus here on the conditions expected to yield extraordinary self-sacrifice, but the complete methods and results are presented in the Supplementary Materials. Consider an infinitely large, Malthusian population of haploid, asexual individuals subdivided into islands of *n* breeders, and let *r* be the consanguinity between two random individuals born on the same island. Breeders produce a large number of offspring and then die. A small fraction *d* of offspring grows wings, while the remaining fraction does not. Thus, dispersal is limited. All offspring then interact in random pairs, giving a fecundity benefit *B* to the recipient at a cost *C* to self. Following this, winged individuals disperse to a random island at a small cost *k*, such that there is a 1–*k* probability of successfully landing on a new island (*16*). Finally, all individuals compete for one of the breeding vacancies with their neighbors on the same island.

To determine the evolutionarily stable (ES) cost-benefit ratio, I calculate the inclusive fitness effect of a mutant actor playing (*B* + *b*,*C* + *c*) for small increments of *b* and *c* and then find the ES marginal cost-benefit ratio, *c*/*b*. Following this, I simulate the effects of dispersal and kin recognition on the ES actual cost-benefit ratio, *C*/*B*, following the tradeoff curve *C* = *B*^*2*^. Extraordinary self-sacrifice evolves under conditions in which *C*/|*B*| > 1 is ES.

In the first model, I consider the effect of actor dispersal. I assume that individuals know whether they will disperse by the presence or absence of wings (*14*). A sedentary actor pays no cost to disperse, and will affect the fitness of neighboring kin of consanguinity *r* who remain on the natal island with probability 1–*d*. Conversely, a dispersing actor successfully migrates to a random island with probability 1–*k*, affecting competition with neighbors of consanguinity 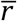. A recipient disperses with probability *d* and so affects neighbors of consanguinity 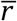 with probability 1–*k*. Otherwise, it remains at home with probability 1–*d* and affects neighbors of consanguinity *r* who also remain on the natal island with probability 1–*d*. With substitution of 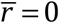, this gives the ES marginal cost-benefit ratio for a sedentary actor

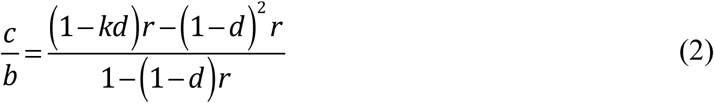

and the ES marginal cost-benefit ratio for a dispersing actor

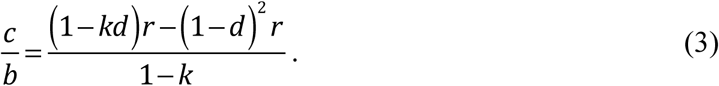

Simulation of these results (Fig. 2) shows that altruism evolves more easily when the actor is sedentary, benefiting its kin, than when the actor disperses and benefits nonkin. Nevertheless, the extent of altruistic behavior in this model is well within ordinary limits. This replicates previous findings by El Mouden and Gardner (*14*).

**Fig. 2.**
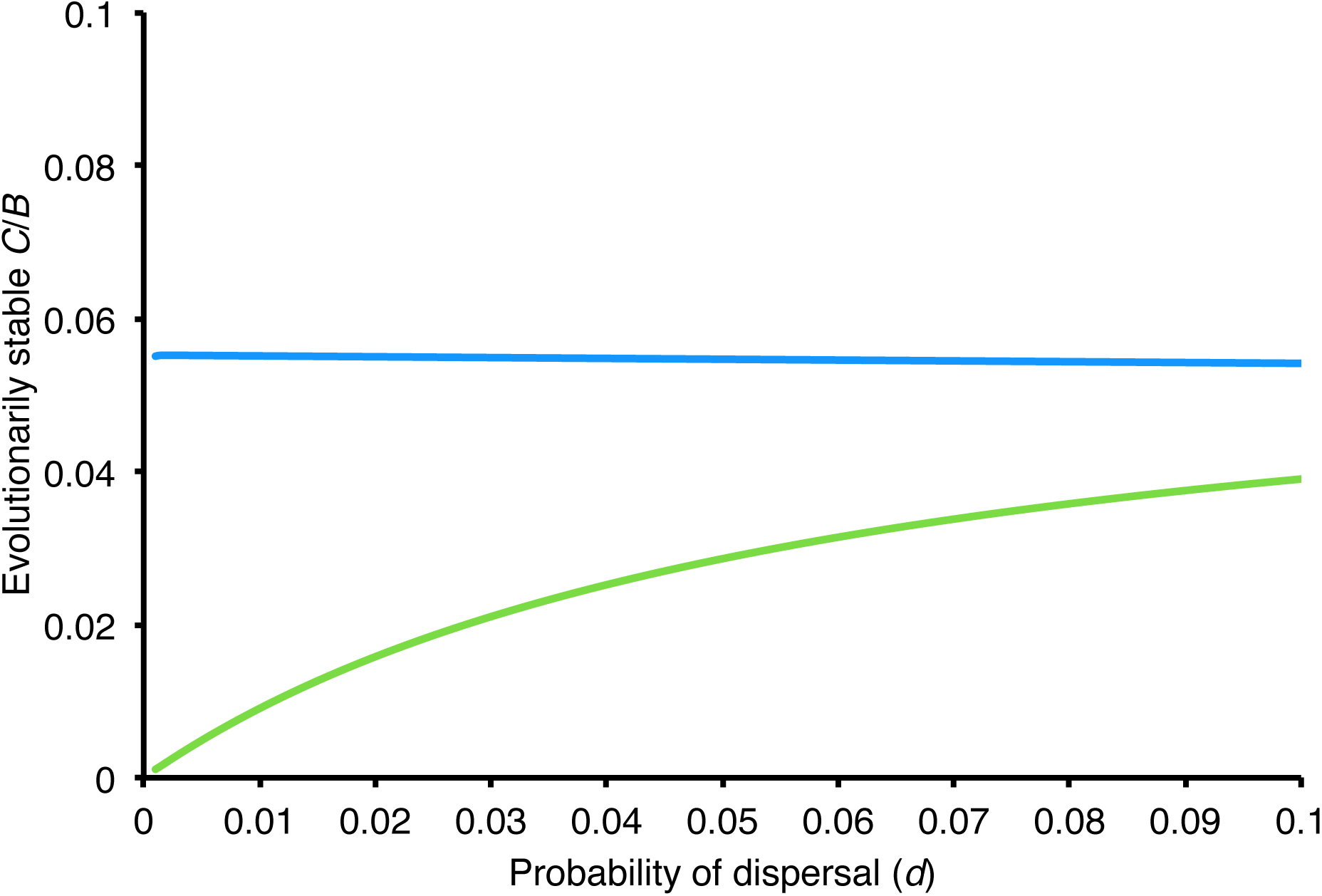
Effect of actor dispersal on the evolution of altruism. This simulation assumes that *n* = 10 and *k* = 0.1, and allows *r* to vary with *d* (see Supplementary Materials). Sedentary actors (blue curve) can sustain a higher cost than can dispersing actors (green curve), especially under limited dispersal. Hence, altruism evolves more easily among sedentary than dispersing actors. However, the scope for altruism in either case remains very small, below the *C*/*B* ≤ 1 limit.

In the second model, I assume that actors are sedentary and instead consider the effect of recipient dispersal. Altruism implies a primary benefit to the recipient, and therefore a secondary cost to the recipient’s neighbors. Consequently, it should be favored when recipients disperse to compete with nonkin. Actors can infer the probability of recipient dispersal in a number of ways; in some cases, actors may even cause the dispersal phenotype by influencing caste determination (*17*). Here, I suppose that actors know whether the recipient is likely to disperse from the presence or absence of wings. This gives the ES marginal cost-benefit ratio for a sedentary actor interacting with a dispersing recipient

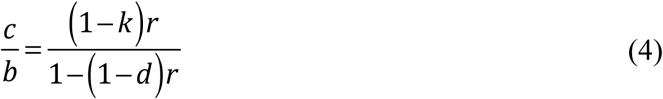

and the ES marginal cost-benefit ratio for a sedentary actor interacting with a sedentary recipient

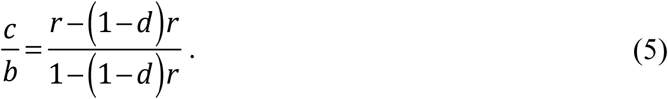

With eq. (4), we now have a case in which extraordinary self-sacrifice can evolve: altruism between a sedentary actor and a dispersing recipient is evolutionarily stable for *C*/|*B*| > 1 (Fig. 3). This is not the case for eq. (5), in which a sedentary actor interacts with a sedentary recipient. This is because the secondary benefit given by the actor’s sacrifice to consanguineous neighbors is offset by the secondary cost imposed on the same neighbors by the recipient’s gain.

**Fig. 3.**
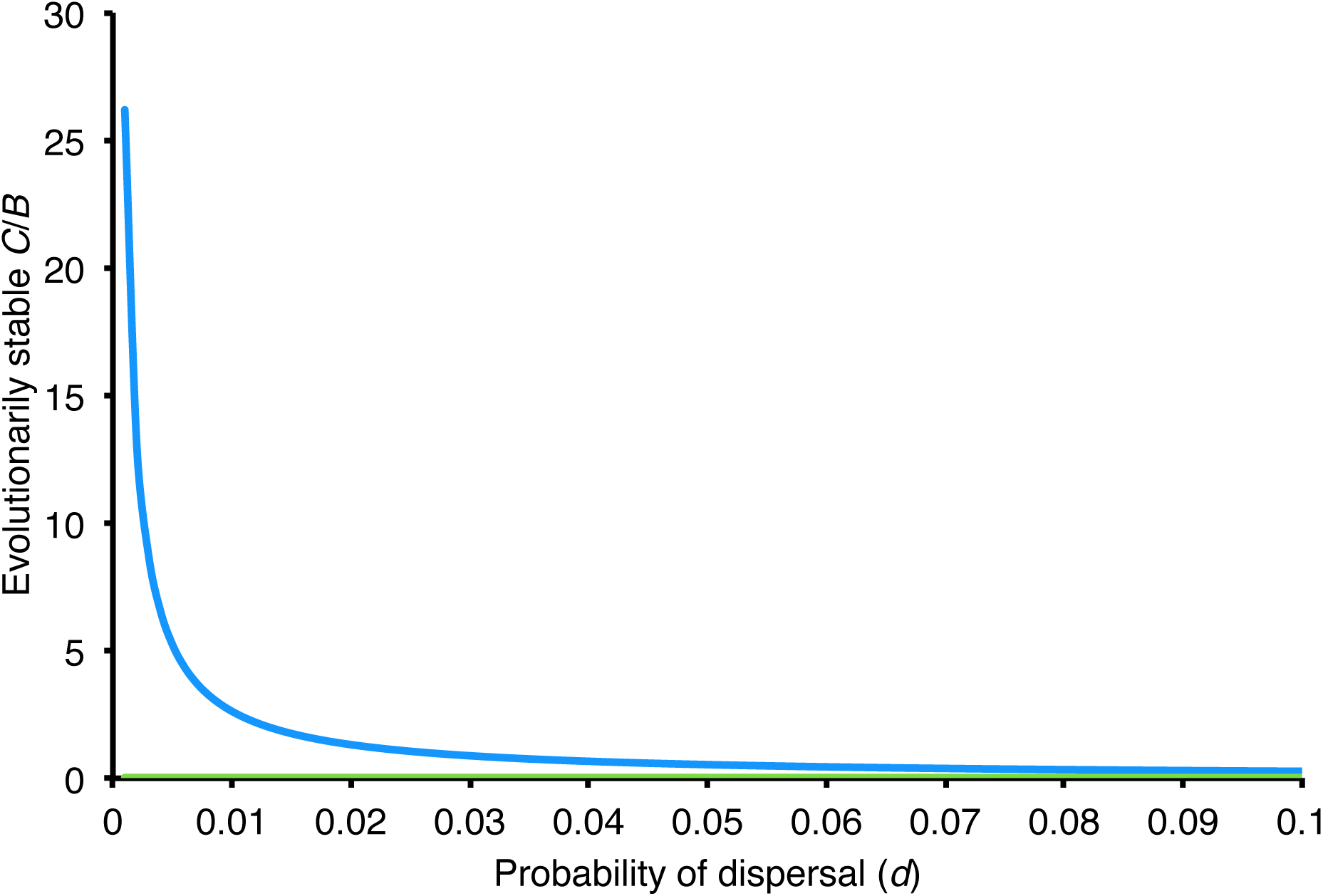
Effect of recipient dispersal on the evolution of altruism. This simulation assumes that *n* = 10 and *k* = 0.1, and allows *r* to vary with *d* (Supplementary Materials). With limited dispersal, sedentary actors interacting with dispersing recipients (blue curve) can sustain much higher costs than can sedentary actors interacting with sedentary recipients (green curve), allowing for the evolution of extraordinary altruism (*C*/*B* > 1).

In the third model, I consider the effect of kin recognition, which allows for the evolution of spite as well as altruism. As in the second model, I assume that actors know whether they and the recipient will disperse by the presence or absence of wings. However, I additionally suppose that offspring learn a song from their parents that originates on the island on which their parents were born (*15*). Thus, all breeders born on the same island sing the same song, but any immigrant breeder sings a different one. All offspring are paired with a random individual drawn from their own island and sing their parent’s song before they interact. It follows that if the actor and recipient sing the same song, then the consanguinity between them is 1, and if they sing different songs, then it is 0.

Two conditions are of particular interest. First, altruism should evolve most easily when the actor is sedentary, the recipient disperses, and the actor and recipient are kin. This is because the action will benefit copies of the focal allele in the recipient as well as in the actor’s neighbors, and it will impose a cost on rival alleles among the recipient’s neighbors. Second, spite should evolve most easily when the actor is sedentary, the recipient sedentary, and the actor and recipient are nonkin. This is because the action will impose a cost on a rival allele borne by the recipient as well as benefit copies of the focal allele in the actor’s neighbors twice over. I find the marginal ES cost-benefit ratio for a sedentary actor interacting with dispersing kin is

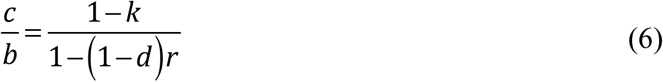

and the marginal ES cost-benefit ratio for a sedentary actor interacting with sedentary nonkin is

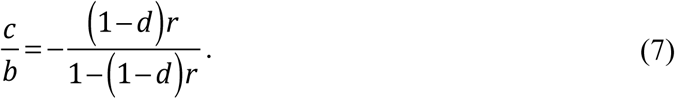

Eqs. (6) and (7) show that both extraordinary altruism and extraordinary spite can evolve (Fig. 4). Thus, kin recognition increases the scope for altruism between sedentary actors and dispersing recipients and increases the scope for spite between sedentary actors and sedentary recipients (*15*).

**Fig. 4.**
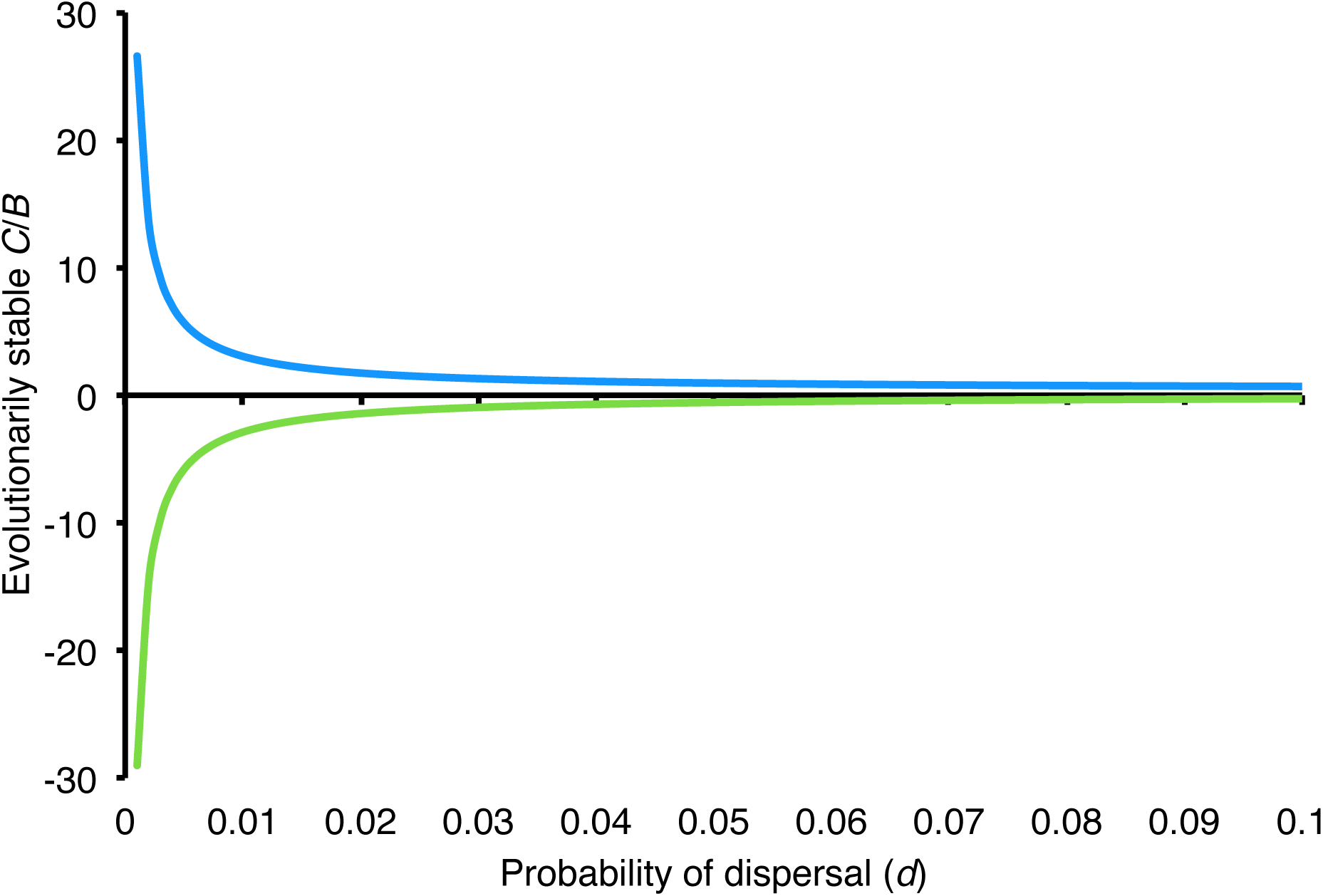
Effect of kin recognition and recipient dispersal on the evolution of altruism. This simulation again assumes that *n* = 10 and *k* = 0.1, and allows *r* to vary with *d* (see Supplementary Materials). With limited dispersal, sedentary actors interacting with dispersing kin (blue curve) can perform extraordinary acts of altruism (*C*/*B* > 1). Likewise, sedentary actors interacting with sedentary nonkin (green curve) can perform extraordinary acts of spite (*C*/*B* < −1).

The models above illustrate the competing effects of dispersal on the evolution of altruism (*10*). On the one hand, dispersing recipients earn a primary benefit and take the secondary costs of altruism elsewhere. On the other hand, dispersal reduces consanguinity between the actor and the recipient, diminishing the primary benefit of altruism, and it reduces consanguinity between the actor and its neighbors, diminishing the secondary benefit of self-sacrifice. In the first model, these effects largely cancel each other out (Fig. 2, blue curve), a finding previously established by Taylor (*8*). In the second model, recipient dispersal is fixed, and so the two negative effects of dispersal on consanguinity become apparent (Fig. 3, blue curve), though they are confounded with one another. Finally, in the third model, both recipient dispersal and actor-recipient consanguinity are fixed, exposing the negative effect of dispersal on actor-neighbor consanguinity (Fig. 4, blue curve).

Taken together, these models show that individuals can evolve to value others’ fitness more than their own. With limited dispersal, extraordinary altruism evolves when sedentary actors interact with dispersing kin and extraordinary spite evolves when sedentary actors interact with sedentary nonkin. From this, we can deduce a simple rule: extraordinary self-sacrifice will evolve when an actor’s neighbors are kin and the recipient’s neighbors are not.

It is reasonable to wonder whether organisms in nature have access to information about kinship and dispersal and make effective use of it. To answer this, first consider the simplest problem: how the actor might know its own probability of dispersal. I assumed above that actors could use the presence or absence of wings to infer dispersal status, which seems reasonable. However, no such inference is actually necessary. Since selection will tend to mold physical and behavioral phenotypes in tandem, we can instead expect sedentary and dispersing individuals to have evolved different inclinations to help, in much the same way that the sexes of a given species may evolve different mating strategies.

The actor’s knowledge of the recipient’s probability of dispersal is a more complicated problem, but not by much. In some cases, actors may themselves determine a recipient’s dispersal type (*17*). In other cases, they may contribute to a resource pool to which dispersing individuals are more inclined or better able to access. Nevertheless, inferential mechanisms are also plausible. Actors can estimate the probability of a recipient’s dispersal with correlates of dispersal status, such as location within the colony, the presence of wings, body size, or odor.

Finally, actors can estimate consanguinity with direct and indirect cues of genealogical kinship (*18*-*19*). For instance, workers may rely on phenotypic cues when interacting with others outside of the colony or at its perimeter. However, they may simply assume that others are kin when operating within the confines of the colony. Hence, kin recognition will depend on context and may consequently produce very different kinds of discrimination.

Eusocial species, which bear many of the most striking adaptations for altruism and spite, confirm the key elements of my argument (*5*, *6*, *17*-*25*). Their colonies consist of close kin, often due to monandry and limited dispersal. Moreover, they comprise a large workforce that produces most of the physical labor and a small number of reproductives that produces most of the offspring. Workers tend to remain with the natal colony whereas reproductives tend to disperse and establish new colonies. Workers and reproductives also have different phenotypes, which implies that they make different decisions and can, in principle, distinguish one another. Finally, kin and nestmate recognition is pervasive, and workers defend the colony from unrelated intruders, often with weapons specifically designed for the task. All of this suggests that the conditions of the theory are both reasonable and reasonably common.

## Acknowledgments

I am grateful to Peter Taylor for feedback on previous drafts of this manuscript and to the attendees of the McMaster University Ecology, Evolution, and Behaviour seminar for comments and discussion.

